# Best templates outperform homology models in predicting the impact of mutations on protein stability

**DOI:** 10.1101/2021.08.26.457758

**Authors:** Marina A. Pak, Dmitry N. Ivankov

**Affiliations:** Center of Life Sciences, Skolkovo Institute of Science and Technology, Moscow 121205, Russia

**Keywords:** FoldX, I-Tasser, AlphaFold, protein stability, homology modeling

## Abstract

**Motivation:** Prediction of protein stability change upon mutation (ΔΔG) is crucial for facilitating protein engineering and understanding of protein folding principles. Robust prediction of protein folding free energy change requires the knowledge of protein three-dimensional (3D) structure. Unfortunately, protein 3D structure is not always available. In this case, one can still predict the protein stability change by constructing a homology model of the protein; however, the accuracy of homology model-based ΔΔG predictions is unknown. The perspectives of using 3D structures of the best templates are also unclear.

**Results:** To investigate these questions, we used the most popular and accurate publicly available tools: FoldX for stability change prediction and I-Tasser for homology modeling. We found that both homology models and best templates worsen the ΔΔG prediction, with best templates performing 1.5 times better than homology models. For AlphaFold models, we also found that the best templates seem to outperform protein models. Our findings imply using the 3D structures of the best templates for ΔΔG prediction if the 3D protein structure is unavailable.

## Introduction

Protein stability is a fundamental property of the protein native three-dimensional (3D) structure. Accurate prediction of folding free energy change upon amino acid mutation facilitates enzyme redesign [1], pathogenicity assessment [2], and optimization of thermostability [3] as well as characterizes our understanding of protein folding principles. Experimental measurements of protein stability change upon mutation are laborious and time-consuming, making computational prediction highly important.

Prediction methods using 3D protein structure produce more robust results than methods using sequence alone [4, 5]. Unfortunately, the 3D structure is unavailable for most of the proteins as seen from a giant and expanding gap between the number of protein sequences and 3D structures [6]. Nevertheless, when the protein 3D structure is unknown, one can still use structure-based prediction methods by building a 3D structure from protein sequence using homology modeling [7] or by highly accurate deep learning algorithm AlphaFold [8]. Homology modeling firstly identifies a homologous protein with the known 3D structure (template) and then builds a 3D protein structure model of the given protein using the identified template. The quality of the constructed model depends on many factors, including template choice and refinement procedure. Obviously, errors in a homology model might influence the prediction of protein stability change. Alternatively, one can use for the predictions the structure of the template directly, without homology model building. As for AlphaFold, at the last stage, it uses constraint relaxation using OpenMM with the Amber99sb force field [8], which could also influence the ΔΔG prediction.

In the present study, we investigated the difference between the usage of homology models and templates in the prediction of protein stability change upon mutation. We used one of the most popular programs for structure-based prediction of protein stability change upon mutation, FoldX [9], and the best publicly available protein structure prediction tool in the web-server CASP category, I-Tasser [10]. We showed that higher accuracy is achieved in the case of utilizing templates in protein stability calculations rather than homology models built from the templates. For the templates with sequence identity greater than 50%, the error of the prediction for homology models is 1.5 times higher than for a template, on average. Moreover, we evaluated performance on multiple-template models of I-Tasser and AlphaFold models from CASP14 and demonstrated that the accuracy of the prediction in this scenario is still worse than for best templates. Our findings strongly suggest using the template structure for the prediction of protein stability change upon mutation if the protein 3D structure is not available.

## 2. Methods

### 2.1 Dataset of protein three-dimensional structures

We compiled a dataset of protein 3D single-domain structures from the Protein Data Bank (PDB) [11] that satisfy the following conditions:

- X-ray determined structures with resolution 2.5 Å or better (required by FoldX [9]);
- Monomeric structures;
- No missing or non-standard residues (required by FoldX [9]);
- Length of 150-250 amino acids. Length restrictions are arbitrarily set to reduce the computational time;
- Globular proteins (according to SCOP [12] classification).

The criteria were submitted in the form of XML representation to RESTful (REpresentational State Transfer) Web Service at the PDB site (http://www.rcsb.org/pdb/software/rest.do; now it is obsolete). To remove too similar proteins in the dataset, we arbitrarily chose the threshold of 90% for the pairwise sequence identities, the distribution is given in Supplementary Figure S1A. The assembled dataset contained 341 entries (Supplementary Table S1).

### 2.2 Template search

To identify the best templates for each protein from the dataset we performed the protein BLAST [13] search against all sequences from PDB:

~~~
blastp -query dataset.faa -out blast.out -db /path/to/pdbaa -outfmt “6 qacc sacc
qseq sseqpident length mismatch gapopen qstart qend sstart send evalue bitscore”
~~~

To eliminate non-homologous proteins, we removed hits having sequence identity less than 30%. We also ignored PDB structures with missing residues. BLAST search found templates for all 341 proteins from the dataset; Supplementary Figure S1B shows the distribution of corresponding sequence identities for the found best templates.

### 2.3 Protein structure modeling by I-Tasser

For homology modeling, we used the stand-alone version of I-Tasser 5.1 [10]. We performed single-template homology modeling by specifying the 3D structure of the protein with the highest sequence identity found at the previous step as a template (-restraint3 flag) and by exclusion of the 3D structure of the query protein (-temp_excl flag):

~~~
runI-TASSER.pl -libdir /path/to/ITLIB/ -seqname $seqname –datadir
/path/to/datadir/$seqname -restraint3 $template:$templatechain -temp_excl
$control:$controlchain -nmodel 1 -runstyle parallel
~~~

For multi-template modeling we excluded the PDB ID of the query protein using -temp_excl flag:

~~~
runI-TASSER.pl -libdir /path/to/ITLIB/ -seqname $seqname –datadir
/ path/to/datadir/$seqname -temp_excl $control:$controlchain -homoflag real
-nmodel 1 -runstyle parallel
~~~

In both cases, we produced one best model for each protein (-nmodel 1).

### 2.4 *In silico* mutagenesis by FoldX

We used FoldX, version 5.1 [9]. First, according to the guidelines of the FoldX manual, we optimized the protein structures by the “RepairPDB” procedure which resolves van der Waals clashes and fixes wrong torsion angles:

~~~
foldx --command=RepairPDB --pdb=file.pdb
~~~

Then, we performed *in silico* mutagenesis by the “BuildModel” tool of FoldX with default settings, the mutation was specified in the individual list mode:

~~~
foldx --command=BuildModel --pdb=file_repaired.pdb
--mutant-file=individual_list.txt
~~~

### 2.5 Dataset of random mutations

For proteins from the dataset, we generated an arbitrarily chosen number of 23 028 random point mutations in positions where both original protein and template had identical amino acid residues. Every such position in original proteins had equal chances to be chosen for *in silico* mutagenesis. After choosing the position, a mutation was chosen randomly from all possible 19 variants with equal probabilities.

### 2.6 Dataset of experimental mutations

We took experimental values of folding free energy changes upon mutation from T1626 dataset [14] which contains high-quality thermodynamic data on single-point mutations. We excluded mutations that contained only melting temperature changes and lacked the values of folding free energy changes. Next, we took PDB entries of the proteins hosting the remaining mutations and for each of them found the best template as described in “Template search” subsection. Then, we selected data in which the mutated position contained the same amino acid residue in the best identified template. The processed dataset resulted in 770 mutations in 33 protein structures (Supplementary Material 2).

### 2.7 AlphaFold models

From all CASP14 targets, we chose those having their 3D structure in PDB with X-ray resolution of 2.5 Å or better and excluded membrane proteins. The resulting two structures are given in Supplementary Table S5. They contained missing residues at the termini, which we ignored (Supplementary Table S5). The PDB structures of the proteins, their best templates (found as described in the “Templates search” subsection), and corresponding AlphaFold models underwent the *in silico* mutagenesis procedure described in the “*In silico* mutagenesis by FoldX” subsection.

### 2.8 Data analysis

Information on solvent accessibility was taken from DSSP [15]. The relative solvent accessibility (*RSA*) of an amino acid residue was calculated according to the equation:

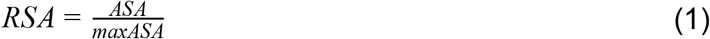

where *ASA* is the solvent accessible surface area and *maxASA* is the maximum possible solvent accessible surface area of the amino acid, both measured in Å^2^.

Following [16] we used the solvent accessibility threshold of 25% to classify residues as exposed or buried.

As a measure of the accuracy of folding free energy change prediction, we used the Pearson correlation coefficient (PCC), *r* :

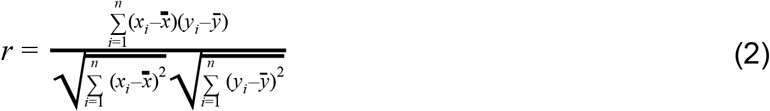

where 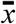 and 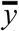 are averages in the *x* and *y* datasets, respectively; *x*_*i*_ and *y*_*i*_ are *i* -th data points from the *x* and *y* datasets.

The standard deviation of PCC was calculated as:

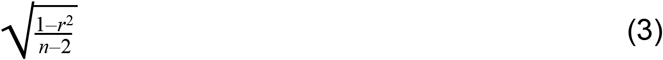

where *r* is PCC, and *n* is the number of considered data points.

Visualization of protein structures was performed in UCSF Chimera [17].

## 3. Results

### 3.1 Design of the computer experiment

To study the influence of homology modeling on protein stability change prediction (further referred to as ΔΔG prediction) we designed a computer experiment represented in Figure 1. In the control part of the experiment, we generated random mutations in the protein 3D structures from the dataset and predicted the ground truth (control) values of the folding free energy change (ΔΔG_control_) associated with the mutations. Next, we emulated the situation when the protein 3D structure is unknown. Specifically, we predicted the effect of the same mutations for the best templates (ΔΔG_template_). Then, we (i) used the structures of the best templates to construct the homology models of the proteins from the dataset and (ii) predicted the effect of the same mutations for the homology models (ΔΔG_homology model_). We considered the drop of agreement between ΔΔG_template_, ΔΔG_homology model_, and ΔΔG_control_ values as an estimate of the effect caused by template or homology model usage in case the 3D structure is unknown. To guarantee the possibility of introducing the same set of mutations into the templates, we generated mutations in the positions where both the control proteins and their best templates had the same amino acid residue.

**Figure 1.**
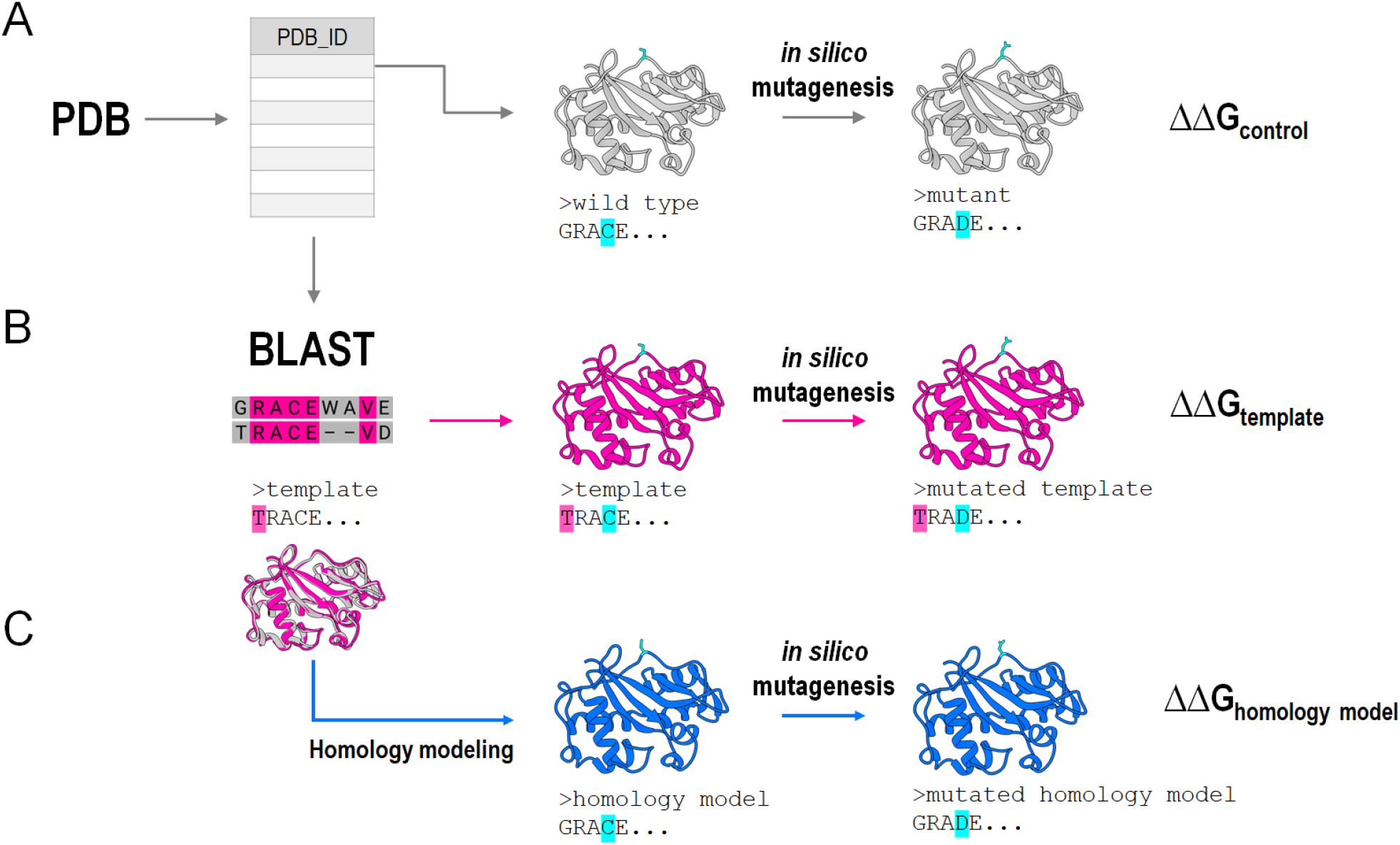
Design of the computer experiment for comparison of approaches to 3D structure unavailability problem. (A) Control experiment: in each protein of the dataset of protein structures we introduced random mutations and predicted their impact on protein stability, obtaining the control ground truth value of ΔΔG (grey structures). (B) For each protein in the dataset, we identified its closest homolog with a known 3D structure and predicted the impact of the same mutations on protein stability. (C) Next, we constructed homology models for each protein based on the best identified homolog and predicted the impact of the same mutations on protein stability as in the control experiment (blue structures).

The programs to perform different steps of the proposed computer experiment are important parameters of the protocol. To predict the effect of mutations on protein stability we chose FoldX, a predictor that manipulates protein 3D structures directly; thus, it is a genuine structure-based tool, which is crucial in the context of the addressed problem. FoldX is widely used and shows one of the best performances among the fast structure-based predictors in the independent tests [18]. To identify the best templates, we used BLAST [13], the classic tool for finding and aligning homologous protein/DNA sequences. As the tool for homology modeling, we utilized I-Tasser, the best template-based method for the automated prediction of protein tertiary structure [10]. I-Tasser server for automated prediction of protein structure has been the best server in The Critical Assessment of protein Structure Prediction (CASP) competition since 2006 [19].

### 3.2 Comparison of the protein stability change prediction for templates, homology models, and control proteins

We calculated the impact of the randomly chosen mutations on protein stability for the original 3D structures (ΔΔG_control_), best templates (ΔΔG_template_), and homology models (ΔΔG_homology model_). We found the correlation with the control experiment to be better for the templates than for homology models, the Pearson correlation coefficient (PCC) being 0.81 ± 0.01 and 0.72 ± 0.01, respectively (Supplementary Figure S2).

Around 10% of points had abnormally high destabilizing values of 10 kcal/mol or higher in either of the calculations. A deeper inspection of these mutations revealed that they correspond to unfavorable conformationally challenging substitutions of small amino acids to bulky ones causing van der Waals clashes (data not shown). The most probable reason is that FoldX does not relax the backbone, so the situation of clashes is not rare. Note that Usmanova *et al*. [20] hypothesized that this FoldX feature is responsible for the anti-symmetry bias in the FoldX predictions. To exclude possible artifacts due to this effect, we trimmed the ΔΔG values using 1.5 IQR and obtained 21014 mutations comprising 91% of the initial dataset. After this removal the correlation with the control experiment remained better for templates than for homology models (Figure 2), PCC being 0.74 ± 0.01 and 0.62 ± 0.01, respectively.

**Figure 2.**
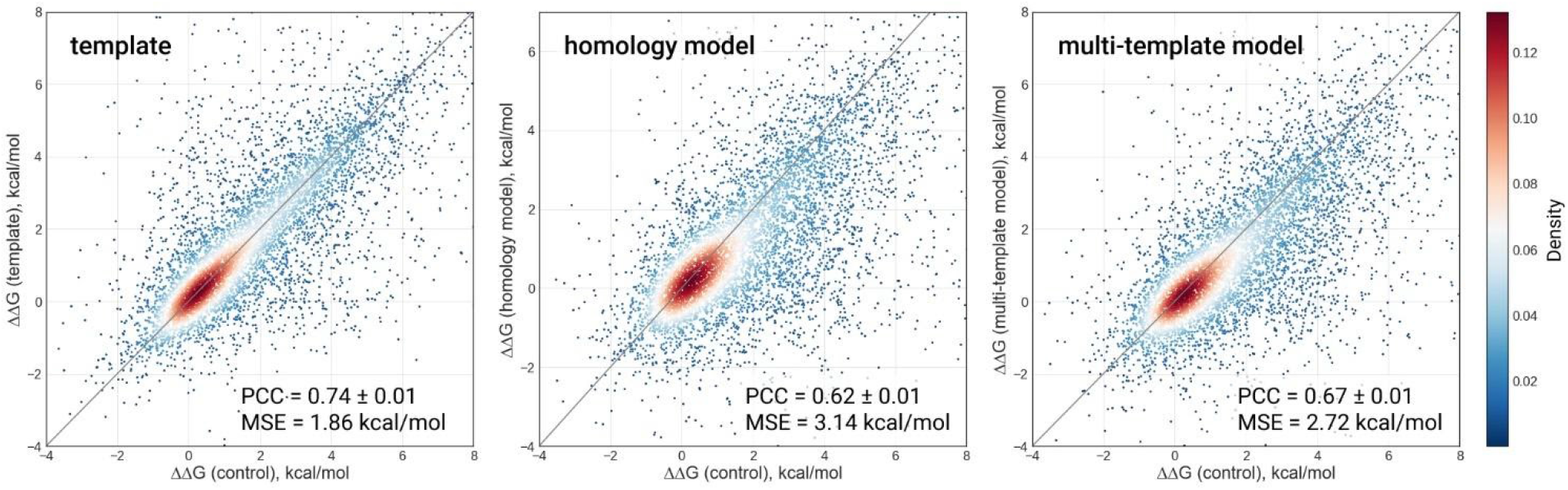
Correlation between the predicted stability changes and control stability changes after removal of points with abnormally high predicted ΔΔG values. PCC between ΔΔG_template_ and ΔΔG_homology model_ is 0.56 ± 0.01, and MSE = 3.49 kcal/mol (plot is not shown). For convenience, we plotted the y = x line in grey.

Interestingly, we observed that the distributions of folding free energy changes for homology models differ (p-value < 0.05) from both the control values and template values (Supplementary Figure S3), while the distributions for template and control values are statistically indistinguishable.

To check how various types of mutations contribute to the observed correlations, we categorized amino acid residues by their charge, hydropathy, polarity, side chain size and relative solvent accessibility (Supplementary Table S2). Then we characterized mutations by the accompanying change of the residue physical properties and calculated PCC within each mutation category (Supplementary Table S3). The correlation with control was always better for templates than for homology models. Thus, the conclusion made above is universal for all mutation categories.

The distributions of the predicted folding free energy changes for template and homology model within each category also remained distinct (Supplementary Table S3). For every category the correlation with control is higher in the case of template, thus, supporting the conclusions above are applicable to every category of mutation.

All things considered, we concluded that in the case of 3D structure unavailability the use of best templates is a more accurate approach for ΔΔG prediction than the use of homology models.

### 3.3 Validation of obtained results on experimental mutations

The ultimate goal of using FoldX or any other prediction method is to accurately estimate experimental values of protein stability change due to mutation. The discrepancy between the predictions made for homology models, the best templates, and the control predictions does not answer the question of how well the ΔΔG predictions agree with experimental data.

To address this question, we repeated the computer experiment for experimentally known mutations. For this purpose, we took T1626, a high-quality dataset of experimental values of folding free energy changes upon single-point mutations [14], filtered it (see Materials and Methods), and processed it as described in the “Design of the computer experiment” section. Figure 3 shows correlations between the predicted and experimental ΔΔG values for original 3D structures, best templates, and homology models for the resulting 752 experimental mutations after removal of outliers.

**Figure 3.**
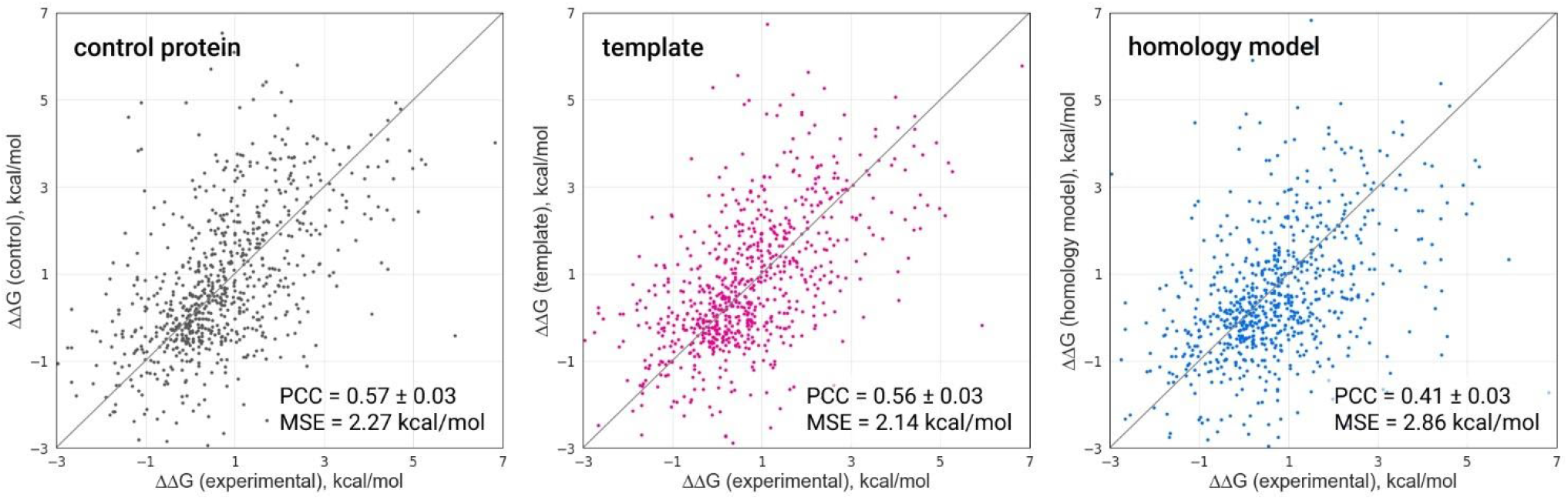
Correlation between the predicted stability changes and the data from the T1626 dataset.

Again, the best templates showed better results than homology models. Moreover, the best templates gave statistically indistinguishable results from the original 3D structures (Figure 3), p-value being 0.49, which probably resulted from the limited amount of data.

For mutations with experimentally measured ΔΔG, PCC between the predictions provided by best templates and original 3D structures (control) was 0.85 ± 0.02, while the predictions for homology models and for original 3D structures (control) correlated at the level of 0.55 ± 0.03 (plots are not shown). This difference is more dramatic than for the randomly chosen mutations (0.74 and 0.62, respectively; see Figure 2) and may be assigned to the different choice of mutations: experimental mutations often contain mutations to alanine or mutations when side-chain is truncated while the dataset of random mutations contains all types of mutations. The PCC between ΔΔG predicted from templates and from homology models is 0.52, which is lower than for randomly generated mutations (0.56, see legend to Figure 2). Due to the limited amount of data, we did not explore different categories of mutations.

It should be noted that the correlation coefficient of 0.57 between experimental free energy changes upon mutation and values obtained for original proteins agrees to a good extent with the PCC = 0.50 obtained for FoldX in the independent comparative assessment of predictors [18].

### 3.4 Dependence of protein stability on sequence identity of the template

The correlations given above characterize the loss of accuracy coupled with using templates and homology models for calculating the impact of a mutation on protein stability. The loss of accuracy is averaged over a wide range of template sequence identities with the proteins at hand (Supplementary Figure S1B). The particular deviation of a prediction from the control value might strongly depend on the sequence identity with the template.

To investigate this question, we additionally constructed the homology models using templates with different sequence identities to the original protein (Figure 4). Similarly to the protocol described above, we introduced random single-point mutations into each homology model and the template of corresponding sequence identity and calculated the correlations with the ground truth for different sequence identities.

**Figure 4.**
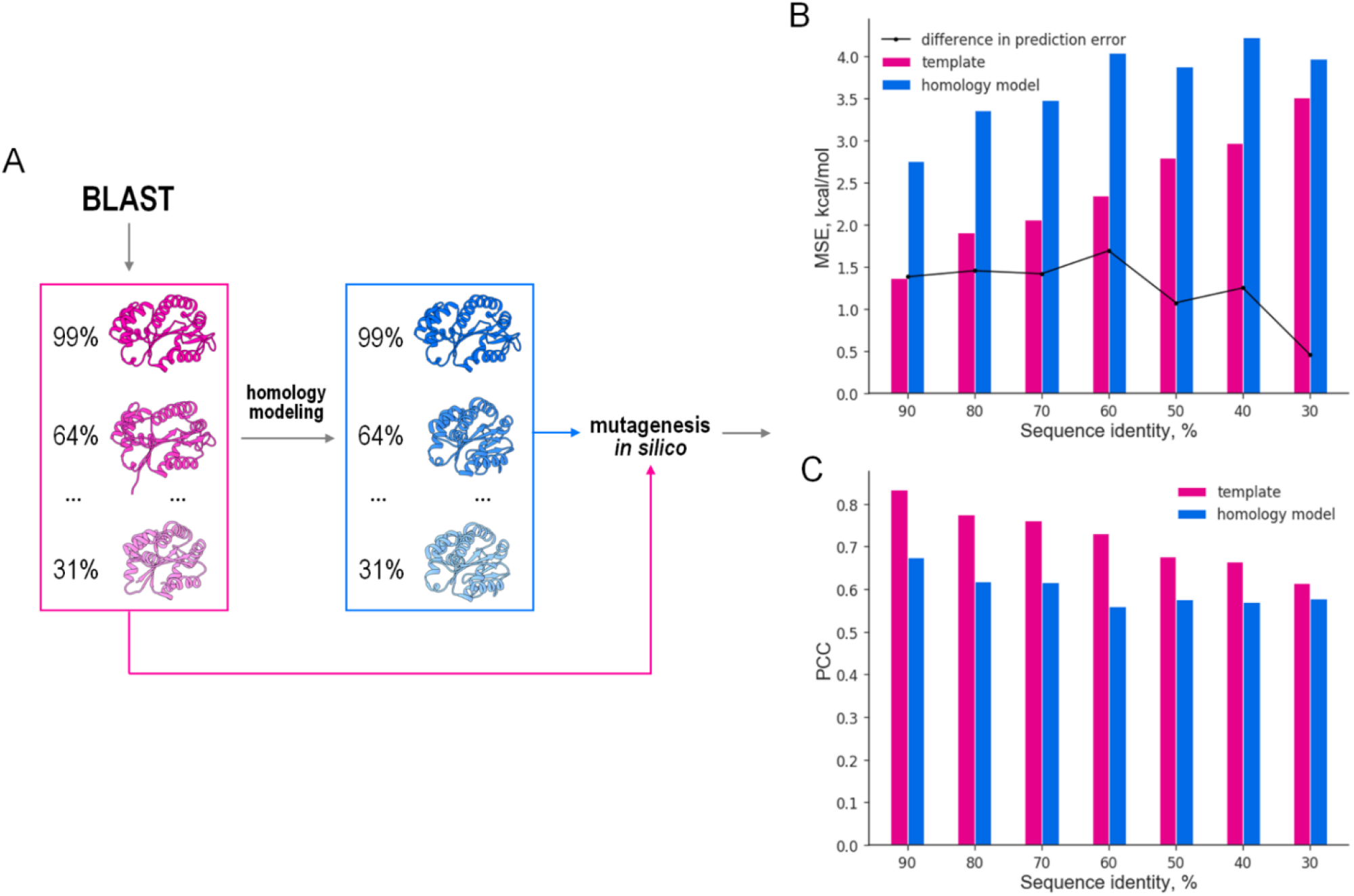
(A) Estimation of dependence on sequence identity. For every protein in the dataset of protein structures, we identified homologous templates with sequence identities in the range of 30% to 100% exclusively. Then we constructed homology models using homologous proteins with different sequence identities as templates. Subsequently, we introduced random mutations in the homology model and the different templates into positions that contain identical residues in the original protein and its templates. (B and C) Finally, for the homologs and homology models, we compared the dependence of the error of prediction (B) and prediction accuracy (C) on the sequence identity of the homolog.

Contrary to the first computer experiment described in Figure 1, where we used only the best template with the highest sequence identity, here we took for each protein a number of different templates with sequence identities in the range from 30% to 100%. Due to computational limitations, we took only 296 original protein structures, with the total number of studied mutations of 3000 and uniform distribution of sequence identities (Figure 4A).

As expected, the accuracy of the prediction dropped with the decrease of sequence identity, while the error of the prediction increased (Figure 4B and 4C). When sequence identity exceeded 50%, the usage of homology models was 1.5 less accurate than that of templates (p-value < 0.05, Mann-Whitney U-test, see also Supplementary Table S4). When approaching the twilight zone of 30% sequence identity the difference in accuracy became less dramatic (Figure 4). Thus, the conclusion of the preferred usage of templates over homology models holds true in the whole range of sequence identities.

### 3.5 Protein stability change prediction with multiple-template models

Using multiple templates in homology modeling improves the model accuracy compared to using just one best template [21]. In I-Tasser, after the alignment of the query sequence to the template structures from PDB, the homology model is constructed through reassembling of structural fragments from multiple templates. In this regard, the one-template restriction for I-Tasser explored above does not reflect the reality, when the multiple-template modeling is the default behavior of I-Tasser.

To explore the perspective of multiple-template homology models for ΔΔG prediction, we constructed the models of original proteins based on multiple templates from PDB using I-Tasser. Then we introduced the same initial set of random mutations and calculated the correlation between folding free energy changes of the models and original proteins. The PCC for multiple-template models is 0.67, which is higher than PCC = 0.62 for homology models but still lower than PCC = 0.74 for the best templates (Figure 2). Again, the distribution of folding free energy changes for multiple-template models is distinct from that for original proteins (p-value of Mann-Whitney U-test < 0.05). Thus, even the homology models obtained in the best multiple-template scenario are worse for ΔΔG prediction than the structures of the best templates.

### 3.6 Protein stability change prediction with AlphaFold models

In the previous section, we explored utilizing multiple templates for model construction. This approach allows addressing the problem of protein structure prediction for the assessment of protein stability change independently from a specific template. This means that we can explore any predicted structure for its perspective for ΔΔG prediction.

Recently, significant advances in protein structure prediction have been accomplished with methods based on convolutional neural networks, such as trRosetta [22] and AlphaFold [8, 23]. These methods are trained on the structures from PDB which, in the context of our study, makes them similar to I-Tasser searching for multiple templates through PDB. At the latest challenge of Critical Assessment of protein Structure Prediction (CASP) AlphaFold remarkably outperformed other methods achieving the Global Distance Test (GDT) score of 90% at which a model might be considered competitive with the experimental structure [24].

Following the logic of the experimental design, we can evaluate the models constructed by AlphaFold in the task of ΔΔG prediction alongside original proteins and templates. Since DeepMind released AlphaFold code to the community, it is tempting to model all the structures used in the presented study and make analysis similar to that done for I-Tasser. This is facilitated by the fact that similar to I-Tasser, AlphaFold offers a convenient option of setting a date and ignoring all the templates from PDB that are newer than that date. However, even with this option, AlphaFold uses weights of the neural network that were obtained for the modern version of PDB so the information about the structure still propagates the predictions. To avoid this inconsistency, we used the models constructed by AlphaFold at CASP14. After filtering (see Methods) it turned out that only two models had templates for corresponding original proteins with the structures available in PDB (Table 1) and, thus, only two models were appropriate for our analysis.

**Table 1.**
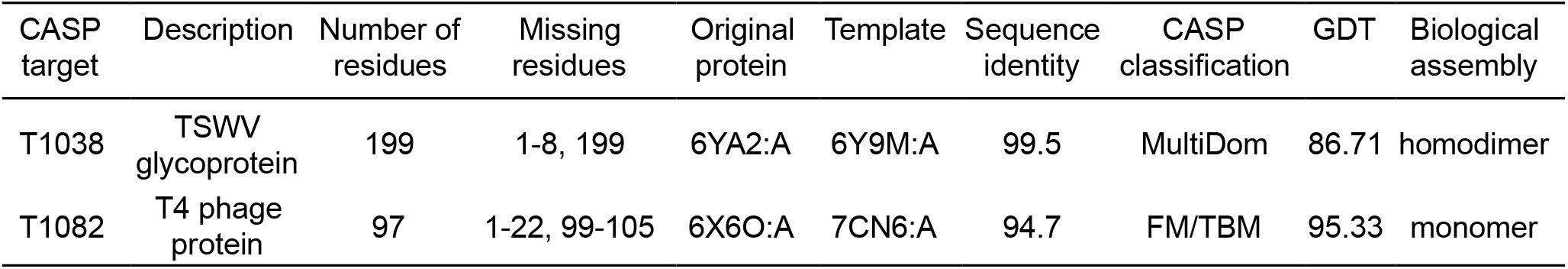
Description of the selected CASP targets and corresponding templates.

| CASP target | Description | Number of residues | Missing residues | Original protein | Template | Sequence identity | CASP classification | GDT | Biological assembly |
| --- | --- | --- | --- | --- | --- | --- | --- | --- | --- |
| T1038 | TSWV glycoprotein | 199 | 1-8, 199 | 6YA2:A | 6Y9M:A | 99.5 | MultiDom | 86.71 | homodimer |
| T1082 | T4 phage protein | 97 | 1-22, 99-105 | 6X6O:A | 7CN6:A | 94.7 | FM/TBM | 95.33 | monomer |

We introduced a set of 1200 random mutations into these two original structures of CASP14 targets, their best templates, and corresponding AlphaFold models. While the correlation coefficients with the control for the best templates and for the AlphaFold models are statistically indistinguishable, the prediction error is greater in the latter case (Figure 5).

**Figure 5.**
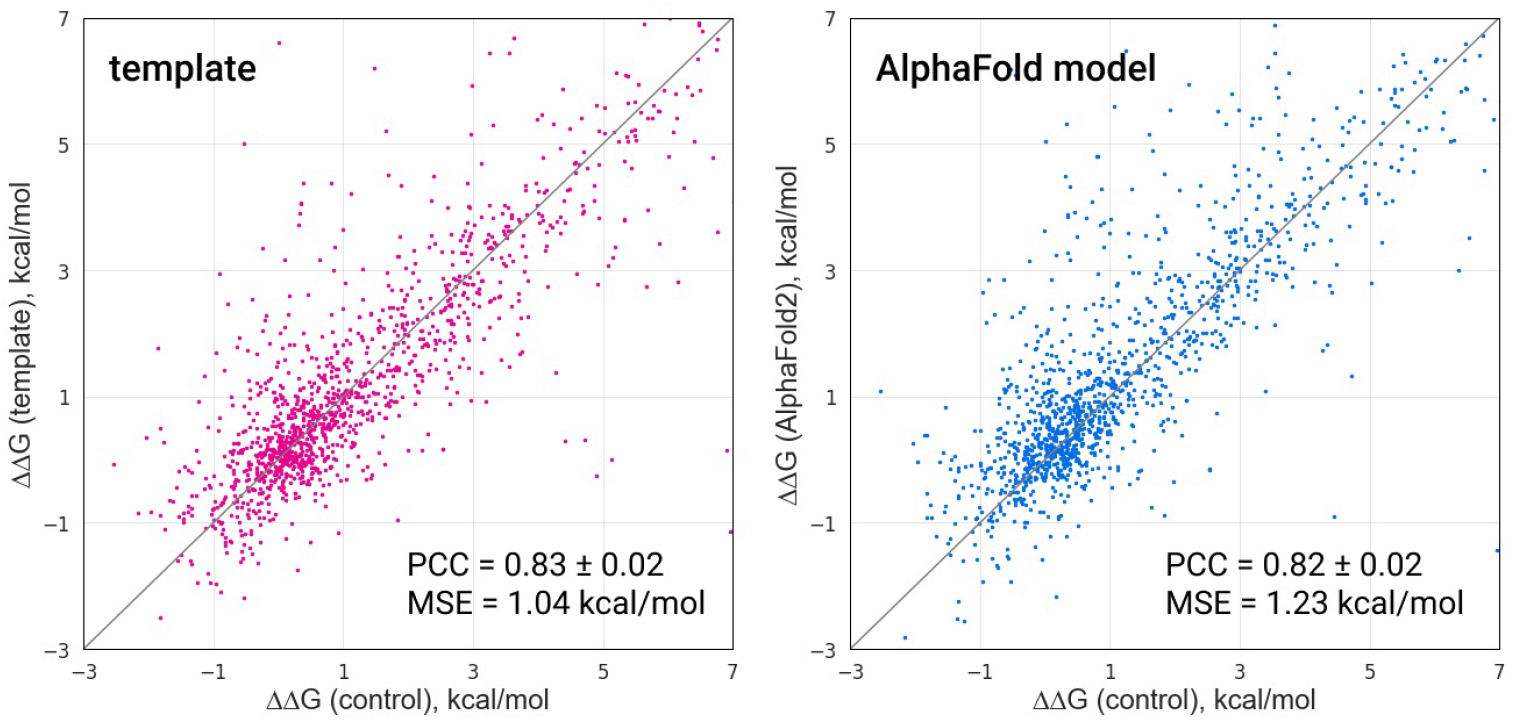
Correlation between control stability changes and predicted stability changes for best templates and AlphaFold models.

As for the distribution of folding free energy changes, they are indistinguishable from original proteins in the case of using templates (Supplementary Table S5, Supplementary Figure S3B).

In surprising consistency with previous findings, the results obtained for AlphaFold models indicate that utilization of predicted protein structure does not improve the accuracy of protein stability change prediction by FoldX. Nevertheless, this conclusion is compromised by the limited amount of structures available for evaluation and as a consequence limited number of introduced random mutations. Thus, structures predicted in the next CASP competitions are desirable to conduct a more comprehensive analysis for AlphaFold.

## 4. Discussion

In the present study, we addressed the problem of 3D structure-based prediction of protein stability change upon mutation when the protein structure is unavailable. We considered a common use-case and chose one of the best, most popular, and user-friendly tools for protein stability prediction, FoldX, and the best publicly available automatic web-server for protein structure construction, I-Tasser. We showed that the use of the structure of the closest homolog is a more accurate approach to ΔΔG prediction than the use of a homology model built from the structure of the closest homolog. The ΔΔG prediction using the closest homolog also better agrees with the experimentally measured protein stability changes than ΔΔG predictions obtained from homology models. Accuracy of ΔΔG prediction using a homology model improves with the sequence identity between the original protein and the template. For the sequence identity of more than 50% one should expect the average error of stability change prediction 1.5 times greater in the case of using a homology model compared to using the closest homolog. Furthermore, the accuracy of ΔΔG prediction is also higher in the case of using a protein structure model constructed based on multiple templates compared to the homology model, but still worse than utilising the closest homolog.

The studied scenario of ΔΔG prediction for a protein without an experimental 3D structure is not purely hypothetical. For example, Pokusaeva *et al*. [25] had to use the homology model of yeast imidazoleglycerol-phosphate dehydratase constructed by I-Tasser to predict the effect of mutations on protein stability. Another example is the ΔΔG predictions made in the homology model myosin VIIa protein constructed by YASARA [26] and followed by 5 ns of molecular dynamics [27]. At the time of those studies, it was unclear to what extent the protein modeling impaired the ΔΔG predictions. Our findings imply that those studies could have been conducted with a higher accuracy if the structure of the template was used, at least for the mutations where the template has the same amino acid residue as the protein at hand.

As for the use of homology modeling for predictions other than ΔΔG, Ittisoponpisan *et al*. [28] published a very good example of the potential use of homology modeling for assessing the pathogenicity of mutations. The authors concluded that missense pathogenic mutations bear structural damage to the protein structure. The authors compared experimental structures and homology models constructed by Phyre2 [29] in the structure-based analysis of the effects of disease-associated mutations. Interestingly, they came to conclusions different from that of the present study: in terms of the TPR/FPR ratio, the results were similar for both experimental structures and homology models, even when the model was built using the template with sequence identity less than 40% [28]. Nevertheless, an important concern here is how well the modeled structure resembles the experimental one in the mutated region. The case studies described by Ittisoponpisan *et al*. [28] illustrate that two homology models built using the templates with the same low sequence identities may have different levels of similarity to the experimental structure in the region of interest and, therefore, produce different results in the analysis of the effect of a mutation. Thus, each situation when the sequence identity of the template is low should be carefully treated individually. It should also be noted that the authors did not assess the performance of the closest homologs or multiple-template protein models in the analysis of the effect of mutations.

Despite the number of ΔΔG predictors is about 30 [30], the analysis presented here covers only FoldX. We chose FoldX for several reasons. First, it has one of the best performances among ΔΔG predictors [18]; the results regarding a method having low accuracy do not have practical reasons. Second, FoldX is a genuinely 3D-structure-based predictor: it changes the mutated amino acids and estimates the effect based on the resulting structure. Oppositely, the machine-learning 3D-structure-based predictors were shown to be insensitive to the details of 3D protein structure, which made Caldararu et al. to question their reliability [31]. As for the predictors from the same category, Eris [32] and Rosetta [33] also show a good performance and have to be tested as well. However, in the present study, we preferred to make an exhaustive analysis of one predictor, FoldX, rather than having significantly less data for the three methods.

More obvious was the choice of a 3D structure predictor. In the web-server CASP category, I-Tasser owns first place in blind predictions of 3D protein structure since 2006. But if to consider all methods, the best choice is obviously AlphaFold since it outperformed other tools at the last two CASP competitions. To study FoldX predictions for AlphaFold models at the same conditions, we had to analyze only two AlphaFold structures from the last CASP round; however, the results seem to agree with the general conclusion made for I-Tasser.

To sum up, we show that if the protein structure is not available it is useless to build the model of protein at hand for ΔΔG prediction, at least by FoldX. Instead, one should utilize the structure of the best template: this will save time on constructing the model and will certainly not decrease the accuracy of the prediction. Surprisingly, the conclusion holds true even for the models made by AlphaFold. We suppose that to make the ΔΔG prediction from protein models relevant, the accuracy should be improved concertedly both for the methods for ΔΔG predictions and for the methods of model construction. In the future, it is most interesting to check systematically the performance of AlphaFold with FoldX, Rosetta, and other ΔΔG predictors using new CASP targets.

## Abbreviations

3D: three-dimensional
DSSP: dictionary of secondary structure for proteins
PDB: protein data bank
SCOP: structural classification of proteins
MW: Mann-Whitney
PCC: Pearson correlation coefficient
MSE: mean standard error
GDT: global distance test
CASP: critical assessment of protein structure prediction
BLAST: basic local alignment search tool
RSA: relative solvent accessibility

## Acknowledgments

We thank Vasily Ramensky for fruitful discussions. The authors acknowledge the usage of the Skoltech HPC clusters Arkuda and Pardus for obtaining the results presented in this paper.

## Notes

### Competing Interest Statement

The authors have declared no competing interest.

