## Supplementary Material 1 for "Best templates outperform homology models in predicting the impact of mutations on protein stability"

**Table S1.** Dataset of control proteins (PDB ID and chain identifier)

| | | | | | | | | | | |
|---|---|---|---|---|---|---|---|---|---|---|
| 1A2Z:A | 1DZR:A | 1GZC:A | 1LBU:A | 1PR9:A | 1UHK:A | 2B0A:A | 2JG6:A | 3F4W:A | 3VTG:A | 4RZ9:A |
| 1A44:A | 1EB6:A | 1H0P:A | 1LHT:A | 1PVH:B | 1UKF:A | 2B82:A | 2O6R:A | 3F9R:A | 3W3E:A | 4TX7:A |
| 1A8L:A | 1EDO:A | 1I76:A | 1LK5:A | 1PVX:A | 1USC:A | 2BEM:A | 2O6S:A | 3GBO:A | 3W42:A | 4U9S:C |
| 1AEC:A | 1EEJ:A | 1I8A:A | 1LQY:A | 1PXV:A | 1UW4:B | 2BK9:A | 2OS3:A | 3GMX:A | 3WYH:A | 4UAV:A |
| 1AGJ:A | 1EJB:A | 1IAM:A | 1M0S:A | 1QB7:A | 1V3W:A | 2BKR:A | 2PCN:A | 3GTT:A | 3ZS3:A | 4WFQ:A |
| 1AH7:A | 1EMY:A | 1ICF:A | 1M2K:A | 1QF9:A | 1V6I:A | 2C4N:A | 2PMK:A | 3H6R:A | 4A02:A | 4XFW:A |
| 1AOE:A | 1ENF:A | 1IFV:A | 1M70:A | 1QFH:A | 1V7R:A | 2CFC:A | 2PMP:A | 3HPE:A | 4A8U:A | 4Z3G:A |
| 1AOL:A | 1EW3:A | 1IIU:A | 1MBS:A | 1QHF:A | 1VAV:A | 2CJL:A | 2PRD:A | 3HZS:A | 4AW8:A | 5AMW:A |
| 1ATG:A | 1EXS:A | 1IO2:A | 1MD6:A | 1QNX:A | 1VDX:A | 2CU9:A | 2PTH:A | 3IOQ:A | 4B96:A | 5B13:B |
| 1AUN:A | 1F2E:A | 1IOO:A | 1MK4:A | 1QR0:A | 1VEC:A | 2E2C:A | 2Q2L:A | 3IP0:A | 4B9C:A | 5B5H:A |
| 1AUO:A | 1F5J:A | 1IQ4:A | 1MNG:A | 1QST:A | 1VMO:A | 2E2D:C | 2Q7A:A | 3JV1:A | 4B9P:A | 5BOW:A |
| 1AVB:A | 1F5V:A | 1IQQ:A | 1MRJ:A | 1QTF:A | 1VS3:A | 2E55:A | 2R9U:A | 3K1R:A | 4BCT:A | 5CFD:B |
| 1AW9:A | 1F99:A | 1IS1:A | 1MTY:D | 1QUA:A | 1WU3:I | 2EBJ:A | 2RI0:A | 3KB5:A | 4DOK:A | 5CFL:A |
| 1BD8:A | 1FCY:A | 1ITV:A | 1MVQ:A | 1QWZ:A | 1X8Q:A | 2ERF:A | 2SCP:A | 3KYF:A | 4EP4:A | 5CPG:A |
| 1BOL:A | 1FNU:A | 1IU8:A | 1N2Z:A | 1R2Q:A | 1XO7:A | 2ET1:A | 2UVD:A | 3LAS:A | 4FRU:A | 5CXK:A |
| 1BSM:A | 1FT5:A | 1IWD:A | 1N81:A | 1R4A:A | 1XS1:A | 2FD5:A | 2VTW:A | 3M6C:A | 4HR6:B | 5DMP:A |
| 1BYI:A | 1FX2:A | 1IYB:A | 1NF9:A | 1RC9:A | 1XYN:A | 2FHE:A | 2WC1:A | 3MND:A | 4J4R:A | 5ECC:A |
| 1C25:A | 1FX4:A | 1JFL:A | 1NFP:A | 1RKU:A | 1Y25:A | 2FJR:A | 2WL1:A | 3MP2:A | 4JRU:A | 5FJQ:A |
| 1CBK:A | 1G12:A | 1JFX:A | 1NIO:A | 1RM8:A | 1Y6H:A | 2FKZ:A | 2X8X:X | 3MW4:A | 4KMY:A | 5FTZ:A |
| 1CFB:A | 1G1T:A | 1JWQ:A | 1NMP:A | 1RPX:A | 1YGH:A | 2GDM:A | 2XB4:A | 3N2N:A | 4L05:A | 5H9I:A |
| 1CFZ:A | 1G3Q:A | 1K12:A | 1NQ7:A | 1RW0:A | 1YOA:A | 2GNU:H | 2Y88:A | 3O7T:A | 4LRU:A | 5HPJ:A |
| 1CHK:A | 1G43:A | 1K3Y:A | 1NQU:A | 1RYP:E | 1YYA:A | 2GPR:A | 2YWX:A | 3OQ3:A | 4NBV:A | 5IC5:A |

| | | | | | | | | | | |
|---|---|---|---|---|---|---|---|---|---|---|
| 1CI3:M | 1G62:A | 1K7C:A | 1NSJ:A | 1S2O:A | 1Z2U:A | 2GSR:A | 2Z3B:A | 3PMD:A | 4NQG:A | 5JJ2:A |
| 1CJW:A | 1G66:A | 1K94:A | 1OA4:A | 1S55:A | 1Z3Q:A | 2GXQ:A | 2ZNR:A | 3PVH:A | 4NUH:A | 5K9A:A |
| 1CZN:A | 1G8A:A | 1KCV:L | 1OAL:A | 1SCZ:A | 1ZIN:A | 2I0W:A | 2ZX2:A | 3QAY:A | 4OY6:A | 5L2V:A |
| 1D2O:A | 1GBG:A | 1KLO:A | 1OKB:A | 1SRD:A | 1ZIR:A | 2IDR:A | 3AJX:A | 3RAB:A | 4PA1:A | 5MRR:A |
| 1DPB:A | 1GBS:A | 1KMV:A | 1OLL:A | 1SUR:A | 2A6Z:A | 2IS9:A | 3BH2:A | 3RBY:A | 4PO4:A | 5OPF:A |
| 1DQN:A | 1GLQ:A | 1KOE:A | 1P2X:A | 1TU7:A | 2AG4:A | 2IWG:B | 3BL6:A | 3SEB:A | 4Q1L:A | 5YH4:A |
| 1DU5:A | 1GNY:A | 1KOP:A | 1P3C:A | 1TYJ:A | 2AGC:A | 2IWX:A | 3CQL:A | 3TOW:A | 4QEC:A | 5YMU:A |
| 1DVO:A | 1GWM:A | 1KS5:A | 1PN9:A | 1U00:A | 2AHN:A | 2J44:A | 3CTK:A | 3TRD:A | 4QMD:A | 6A4F:A |
| 1DXJ:A | 1GWY:A | 1L8F:A | 1PPO:A | 1UCD:A | 2AYH:A | 2JA9:A | 3E9U:A | 3UFC:X | 4QT4:A | 6EDG:A |

**Table S2.** Residue properties

| Property | Type | Residues |
|---|---|---|
| Charge | Positive | R, H, K |
| | Neutral | A, N, C, Q, G, I, L, M, F, P, S, T, W, Y, V |
| | Negative | D, E |
| Hydropathy | Hydrophobic | A, C, I, L, M, F, W, V |
| | Neutral | G, H, P, S, T, Y |
| | Hydrophilic | R, N, D, Q, E, K |
| Polarity | Polar | R, N, D, Q, E, H, K, S, T, Y |
| | Nonpolar | A, C, G, I, L, M, F, P, W, V |
| Side chain size | Very small | A, G, S |
| | Small | N, D, C, P, T |
| | Medium | Q, E, H, V |
| | Large | R, I, L, K, M |
| | Very large | F, W, Y |
| Solvent accessibility | Buried (RSA < 25%) | |
| | Exposed (RSA ≥ 25%) | |

**Table S3.** Comparison of correlation coefficients for different categories of mutations

| **Type of change** | **Percent of mutations of the type** | **PCC (homology model)** | **PCC (template)** |
|---|---|---|---|
| **Polarity** | | | |
| *Change of polarity* | | | |
| polar to nonpolar | 25 | 0.53 ± 0.01 | 0.737 ± 0.008 |
| nonpolar to polar | 24 | 0.59 ± 0.01 | 0.748 ± 0.008 |
| *No change in polarity* | | | |
| nonpolar to nonpolar | 22 | 0.57 ± 0.01 | 0.773 ± 0.008 |
| polar to polar | 22 | 0.52 ± 0.01 | 0.758 ± 0.008 |

| | | | |
|---|---|---|---|
| **Hydropathy** | | | |
| *Change in hydropathy* | | | |
| hydrophobic to hydrophilic | 11 | 0.65 ± 0.01 | 0.79 ± 0.01 |
| hydrophilic to hydrophobic | 13 | 0.39 ± 0.01 | 0.64 ± 0.01 |
| neutral to hydrophilic | 9 | 0.58 ± 0.02 | 0.77 ± 0.01 |
| hydrophilic to neutral | 9 | 0.53 ± 0.02 | 0.74 ± 0.01 |
| neutral to hydrophobic | 12 | 0.56 ± 0.01 | 0.75 ± 0.01 |
| hydrophobic to neutral | 10 | 0.60 ± 0.02 | 0.79 ± 0.01 |
| *No change in hydropathy* | | | |
| neutral to neutral | 7 | 0.55 ± 0.02 | 0.75 ± 0.02 |
| hydrophobic to hydrophobic | 13 | 0.51 ± 0.01 | 0.74 ± 0.01 |
| hydrophilic to hydrophilic | 8 | 0.49 ± 0.02 | 0.71 ± 0.01 |
| **Charge** | | | |
| *Change to the opposite charge* | | | |
| negative to positive (-+) | 2 | 0.56 ± 0.04 | 0.72 ± 0.03 |
| positive to negative (+-) | 1 | 0.47 ± 0.04 | 0.75 ± 0.03 |
| *Loss of charge* | | | |
| positive to neutral (+0) | 10 | 0.42 ± 0.02 | 0.74 ± 0.01 |
| negative to neutral (-0) | 9 | 0.48 ± 0.02 | 0.71 ± 0.01 |
| *No change in charge* | | | |
| neutral to neutral (00) | 51 | 0.608 ± 0.007 | 0.786 ± 0.005 |
| positive to positive (++) | 1 | 0.22 ± 0.05 | 0.64 ± 0.04 |
| negative to negative (- -) | 1 | 0.39 ± 0.07 | 0.67 ± 0.05 |
| *Charge acquirement* | | | |
| neutral to positive | 10 | 0.62 ± 0.02 | 0.75 ± 0.01 |
| neutral to negative | 8 | 0.72 ± 0.02 | 0.81 ± 0.01 |
| **Side chain size** | | | |
| very large change, from larger residue (-4) | 2 | 0.55 ± 0.03 | 0.67 ± 0.09 |
| large change, from large residue (-3) | 8 | 0.56 ± 0.01 | 0.70 ± 0.01 |
| medium change, from large residue (-2) | 12 | 0.55 ± 0.01 | 0.73 ± 0.01 |
| small change, from large residue (-1) | 17 | 0.59 ± 0.01 | 0.73 ± 0.01 |
| no change (0) | 16 | 0.62 ± 0.01 | 0.804 ± 0.009 |
| small change, from smaller residue (1) | 17 | 0.58 ± 0.01 | 0.802 ± 0.009 |
| medium change, from smaller residue (2) | 12 | 0.66 ± 0.01 | 0.82 ± 0.01 |
| large change, from smaller residue (3) | 7 | 0.62 ± 0.01 | 0.81 ± 0.01 |
| very large change, from smaller residue (4) | 2 | 0.42 ± 0.04 | 0.74 ± 0.03 |
| **Relative solvent accessibility of the mutated residue** | | | |
| Exposed | 40 | 0.633 ± 0.007 | 0.713 ± 0.006 |
| Buried | 52 | 0.570 ± 0.007 | 0.767 ± 0.005 |

**Table S4.** Comparison of protein stability change prediction using templates and using homology models for different sequence identities

| Sequence identity, % | Templates | | | Homology models | | | Difference in prediction error, kcal/mol |
|---|---|---|---|---|---|---|---|
| | PCC | MSE, kcal/mol | p-value (MW U-test) | PCC | MSE, kcal/mol | p-value (MW U-test) | |
| 30 | 0.614 ± 0.014 | 3.515 | 0.13 | 0.578 ± 0.015 | 3.975 | < 0.05 | 0.460 |
| 40 | 0.665 ± 0.014 | 2.975 | 0.02 | 0.570 ± 0.015 | 4.227 | < 0.05 | 1.252 |
| 50 | 0.676 ± 0.013 | 2.801 | 0.09 | 0.577 ± 0.015 | 3.877 | < 0.05 | 1.076 |
| 60 | 0.731 ± 0.012 | 2.345 | 0.19 | 0.560 ± 0.015 | 4.038 | < 0.05 | 1.693 |
| 70 | 0.761 ± 0.012 | 2.058 | 0.08 | 0.616 ± 0.014 | 3.478 | < 0.05 | 1.420 |
| 80 | 0.775 ± 0.012 | 1.905 | 0.18 | 0.618 ± 0.014 | 3.361 | < 0.05 | 1.456 |
| 90 | 0.834 ± 0.010 | 1.370 | 0.19 | 0.674 ± 0.013 | 2.754 | < 0.05 | 1.384 |

**Table S5.** Comparison of protein stability change prediction using templates and using AlphaFold2 models

| Dataset | Templates | | | AlphaFold2 models | | |
|---|---|---|---|---|---|---|
| | PCC | MSE, kcal/mol | p-value (MW U-test) | PCC | MSE, kcal/mol | p-value (MW U -test) |
| All | 0.826 ± 0.016 | 1.043 | 0.15 | 0.818 ± 0.016 | 1.232 | < 0.05 |
| 6YA2 | 0.817 ± 0.025 | 0.990 | 0.08 | 0.819 ± 0.025 | 1.113 | < 0.05 |
| 6X6O | 0.842 ± 0.022 | 0.966 | 0.07 | 0.844 ± 0.022 | 1.080 | < 0.05 |

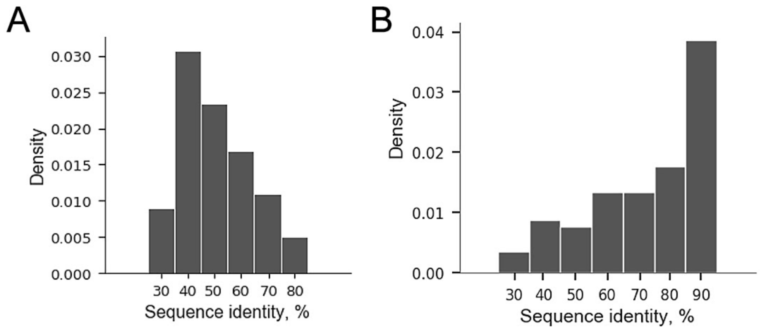


**Figure S1.** (A) Distribution of pairwise sequence identities within the dataset of original proteins (A) and among the best templates (B)

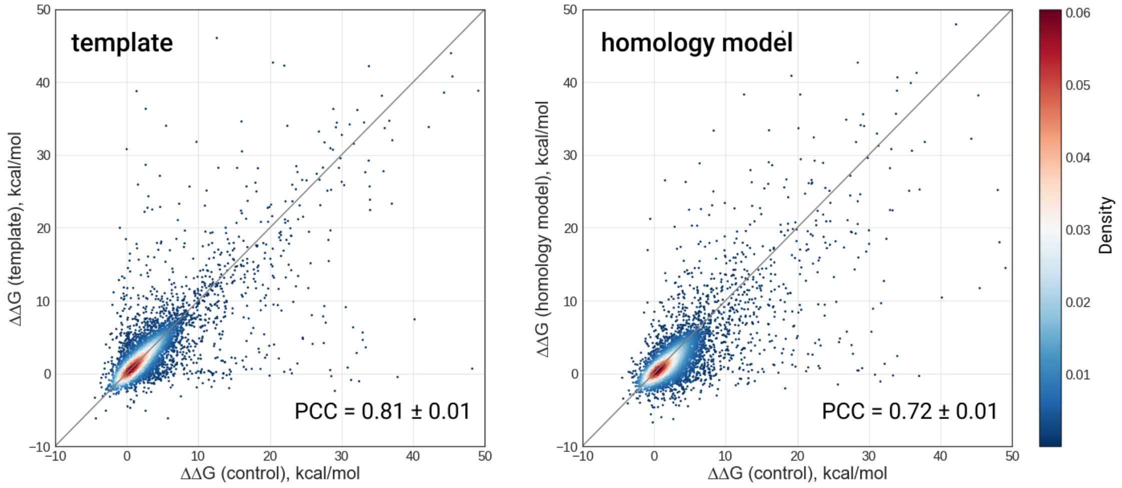


**Figure S2.** Correlation between the predicted stability changes for templates (left) and homology models (right) vs. predicted stability changes of original proteins

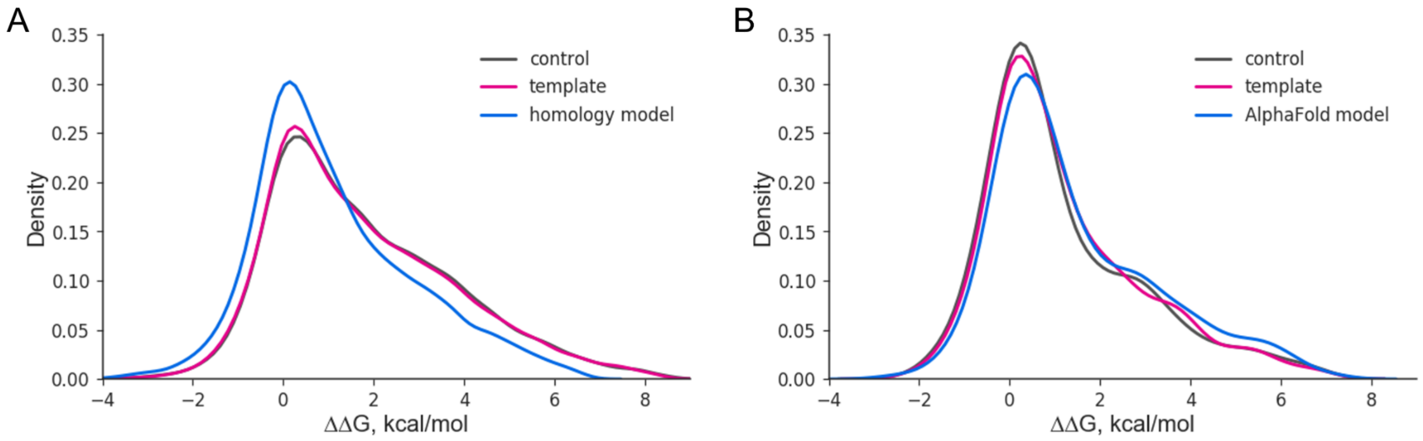


**Figure S3.** Distribution of calculated folding free energy changes for homology models (A) and AlphaFold models (B). using Kruskal-Wallis test. The test indicated that the distributions within a category are distinct (p-value < 0.05), in consequence we estimated the correlations of folding free energy changes with control for template and homology model within each category.
